# A positive list administered immunity system restricts the body size of insects

**DOI:** 10.1101/2024.06.30.600924

**Authors:** Norichika Ogata, Tomoko Matsuda

**Affiliations:** Nihon BioData Corporation, Kawasaki, Kanagawa, Japan; Graduate school of Engineering, Osaka University, Yamadaoka, Osaka, Japan

## Abstract

There are several explanations for the extinction of ancient giant insects. Here, we present a new hypothesis suggesting that the innate immune system limits the body sizes of insects. In this study, we co-cultured bacteria, fungi, and insect blood cells, and performed single-cell RNA sequencing analyses of insect blood cells to determine their division of labor. In the innate immune system, prohibited molecules are listed as signals of invasion. The increasing diversity of organisms makes this list extensive. The burden of managing such an extensive list leads to a division of labor among blood cells and reduces the effective number of blood cells. Our simulation indicates that a reduced number of effective blood cells cannot protect a giant body from invaders.

## Introduction

In general, countries administer their military forces using negative lists. Negative list operations, which allow military forces to do anything except what is explicitly prohibited, give them more flexibility and power. Military forces face complex and various situations and have missions that must be completed. However, the Japanese Self-Defense Forces (Jiei-tai) are administered using a positive list. A study comparing negative list-administered military forces and positive list-administered Jiei-tai in the context of the marginal cost pricing principle in public economics indicated that adopting a positive list operation is a proactive method of concentrating the resources of the authorized organization in areas that will be of great benefit to the entruster, without forcing the Jiei-tai to perform unnecessary tasks (Okuda 2018). Since the acquired immune system is a negative list type immune system and the innate immune system is a positive list type immune system, the immune system of insects that do not have acquired immunity may achieve resource concentration. In this context, insects are possibly thought to be reducing certain aspects; concentrating resources is inextricably linked to cutting out areas that were not selected. In this study, we hypothesized that insects reduce their body size and conducted simulations regarding body size and the number of effective immune cells. In insect immune systems, blood cells are the main component, as demonstrated by a study using Coleopteran larvae (Haine et al. 2008). To estimate the number of effective immune cells, we performed single-cell RNA sequencing of Coleopteran insect larvae and estimated the size of the positive list. We determined that there is a division of labor among blood cells by observing the frequency of adhesion in co-cultures of bacteria, fungi, and Coleopteran insect blood cells.

## Materials & Methods

We obtained *Coleopteran* insect larvae (*Trypoxylus dichotomus*) from the Seven Face Mountain, Kawasaki, Japan. We aseptically dissected the larvae as previously described (Ogata and Iwabuchi 2017) and cultured midgut contents in Shields and Sang M3 Insect Medium with 10% Fetal bovine serum. We isolated a gram-positive bacteria, *Enterococcus sp*. (genome sequencing data is deposited as NCBI SRA DRR142835) and a fungus, *Mucor sp*. (genome sequencing data is deposited as NCBI SRA DRR142836). These microorganisms were continuously subcultured in the M3 media. We aseptically obtained larval blood cells and cultured them in the M3 media (25 C degree). Co-cultures of bacteria, fungus, and blood cells were performed in the M3 media. We observed 135 blood cells using a microscope as previously described (Ogata and Iwabuchi 2017). We noted the frequency of insect blood cellular adhesion to bacteria and fungus. The adhesion frequency was statistically tested using Fisher’s exact test. Aseptically obtained 92 larval blood cells were isolated and sequenced using Fluidigm C1 and Illumina NGS (single cell RNA sequencing data is deposited as NCBI SRA PRJDB6473). Sequence data were mapped to the genome as previously described (Ogata, Yokoyama, and Iwabuchi 2012; Ogata 2021) and we plotted the expression of a hemocytin-like gene (IABQ01002019.1, Locus 703) and a peptidoglycan recognition protein (PGRP)-like gene (IABQ01000018.1 - IABQ01000154.1, Locus 8) of each cell. In the simulation, we simulated a 0.006 mm^3^ space (1000 *μ*m × 600 *μ*m × 10 *μ*m). We simulated the 480 invaders randomly in the space according to the previous study (Haine et al. 2008) and 20, 60 and 180 blood cells according to the previous studies (2,000 - 60,000 blood cells per 1 *μ*L(mm^3^) is general) (Belal and Gad 2023; Lubawy and Slocinska 2020; Kong et al. 2018; Vivekanandhan et al. 2022; Gao et al. 2022). The simulated blood cell moved 10*μ*m per 15 seconds. When the simulated blood cells encountered the simulated invaders, the blood cell kills the invader. These simulations were performed using Lisp (GNU CLISP 2.49.92) (Barski 2010).

## Results and Discussion

We observed 135 blood cells in co-cultures of bacteria, fungi, and Coleopteran insect blood cells. Among these, one blood cell adhered to both bacteria and fungi, 10 cells adhered only to fungi, 53 cells adhered only to bacteria, and 70 cells did not adhere to anything (Figure 1). These results indicate a division of labor among blood cells (p = 0.049). We also analyzed the gene expressions of 92 blood cells, focusing on a hemocytin-like gene and a peptidoglycan recognition protein (PGRP)-like gene (Figure 2). The expression of the hemocytin-like gene and the PGRP-like gene was mutually exclusive (p = 9.6e^−6^) in a regression analysis. These findings suggest that the division of labor among blood cells is rooted in their gene expression. This division of labor implies that the number of effective immune cells is smaller than the total hemocyte count observed in previous studies.

**Figure 1.**
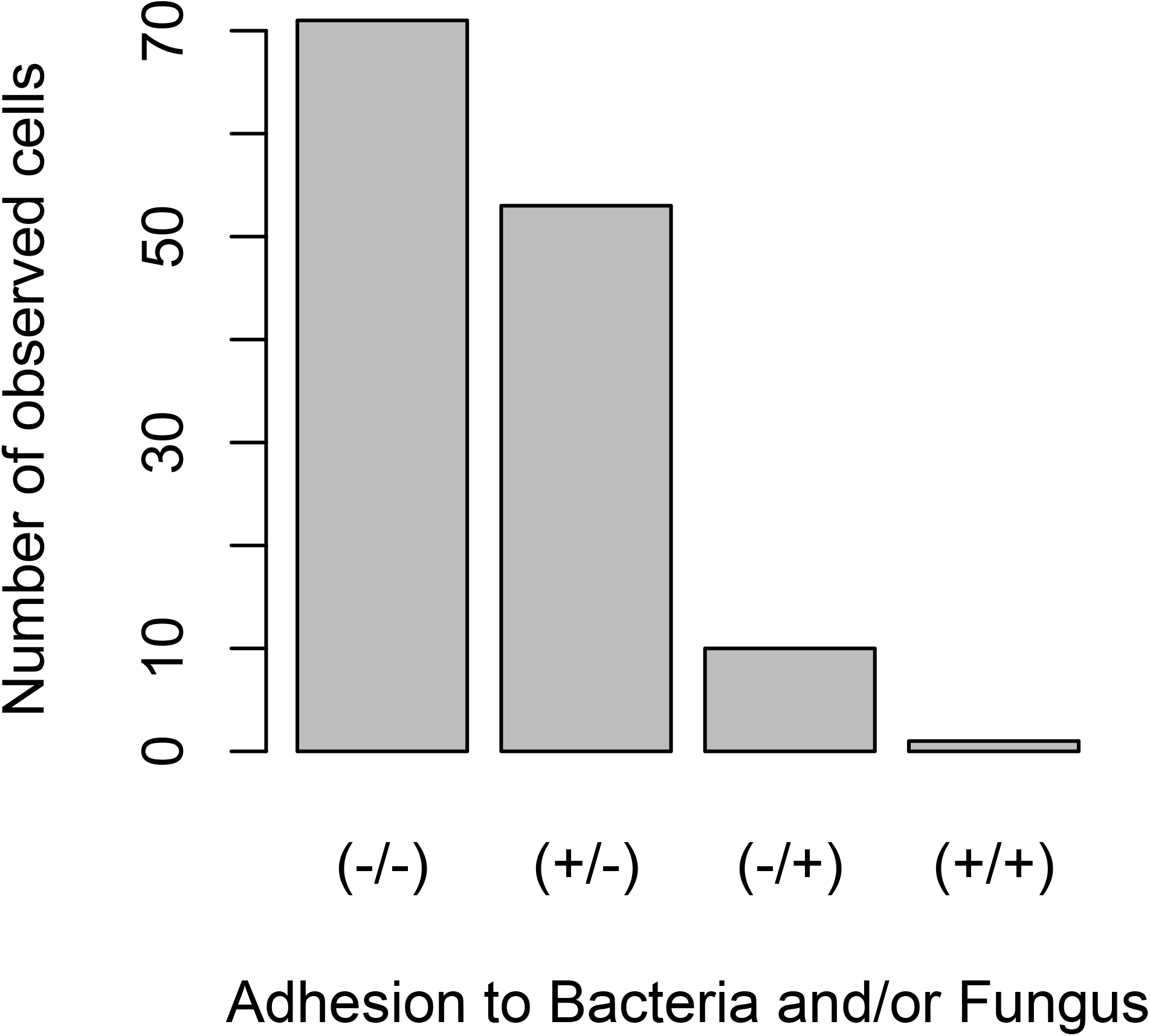
Co-cultures of bacteria, fungi, and insect blood cells. We observed the blood cells and recorded their adherence patterns. (−/−) indicates cells that did not adhere to anything. (+/−) indicates cells that adhered only to bacteria. (−/+) indicates cells that adhered only to fungi. (+/+) indicates cells that adhered to both bacteria and fungi.

**Figure 2.**
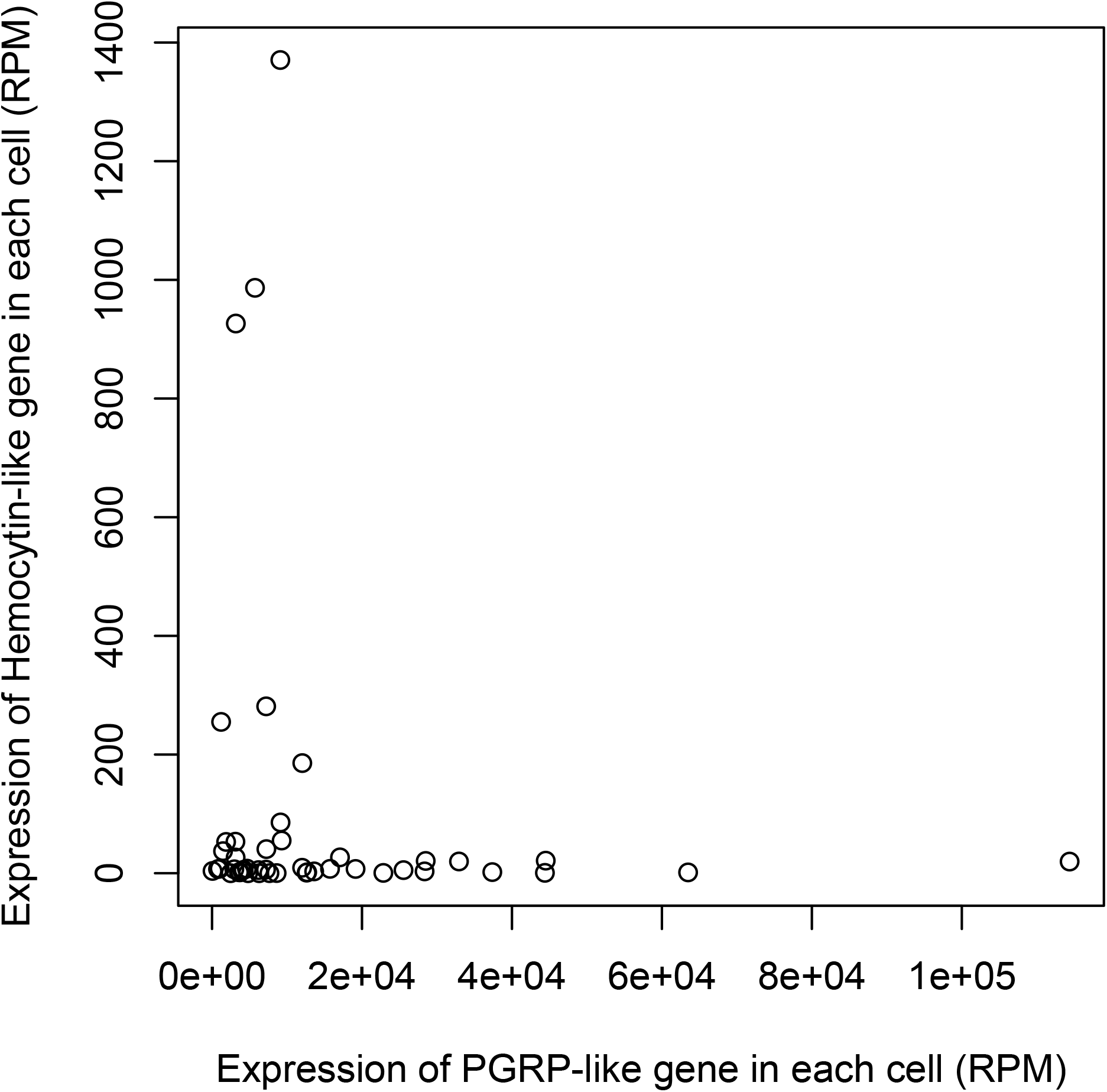
Comparison of PGRP-like Gene Expression and Hemocytin-like Gene Expression in Each Cell. We performed single-cell RNA sequencing analyses of 92 insect cells. The expression levels of the PGRP-like gene and the Hemocytin-like gene were plotted for each cell.

**Figure 3.**
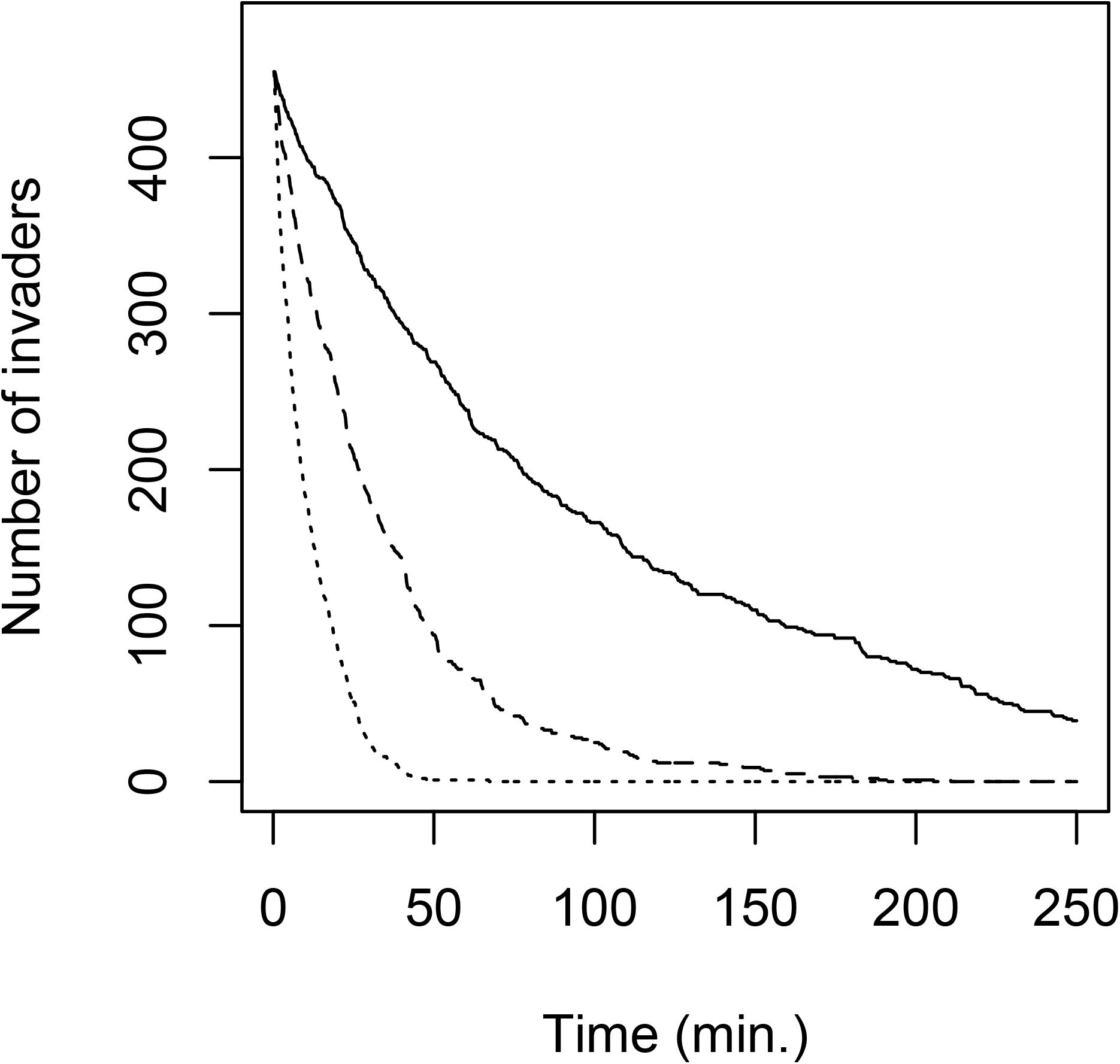
Results of the Blood Cell Killing Invaders Simulation. The solid line represents the number of invaders with 20 blood cells. The dashed line represents the number of invaders with 60 blood cells. The dotted line represents the number of invaders with 180 blood cells.

We simulated the process of removing invaders. With 180 and 60 simulated blood cells, all invaders were eliminated in 30 and 210 minutes, respectively. In the case of fast-growing invaders like *Vibrio parahaemolyticus*, the number of invaders increased by only 8 after 30 minutes and by 16,000 after 120 minutes. The reduction in effective immune cells due to the division of labor among blood cells is critical in larger body sizes. The risk of failing to detect an invader increases with the volume of the body cavity. For instance, assuming a 0.1% chance of failing to remove microorganisms in the 0.006 mm^3^ space we previously defined, the probability of not missing microorganisms is 0.88 in the entire body fluid up to 4 ml, obtainable by squeezing 35 g of beetle larva. However, if the weight increases to about 200 g and body fluids increase approximately fivefold, the same probability decreases to 0.20. If even one of the small compartments discussed in the previous simulation were to fail, it could lead to eventual erosion of the entire open vasculature and spacious hemocoel. In essence, maintaining a large body size while effectively combating a variety of invaders with a positive list proves challenging.

Traditionally, the extinction of ancient giant insects has been attributed to declining atmospheric oxygen levels. However, accumulating evidence challenges this theory. Modern insects maintain very low blood oxygen levels (11 *μ*mol/L, (Stief and Eller 2006)), and insect organs moved to a cultured environment showed decreased expression of hypoxia-related genes (Ogata, Yokoyama, and Iwabuchi 2012). Furthermore, the discovery of insect lungs, composed of trachea and blood cells, has been reported (Locke 1997). A recent study suggested that insect blood cells can transport oxygen (Shin et al. 2024). A study suggested that birds were responsible for the extinction of ancient giant insects (Clapham and Karr 2012). Similarly, our study proposes that predators were a significant factor in the extinction of ancient giant insects.

## Conclusions

In this study, we have proposed a novel explanation: that the innate immune system restricts the body sizes of insects through co-cultures, single-cell RNA sequencing, and simulations.

